# Adaptive switch to sexually dimorphic movements by partner-seeking termites

**DOI:** 10.1101/340919

**Authors:** Nobuaki Mizumoto, Shigeto Dobata

**Affiliations:** Laboratory of Insect Ecology, Graduate School of Agriculture, Kyoto University, Kitashirakawa-oiwakecho, Sakyo-ku, Kyoto, Japan 606-8502

**Keywords:** Random walk, Tandem runs, Sexual selection, Social insects, Movement ecology

## Abstract

When searching for targets whose location is not known, animals should benefit by adopting movement patterns that promote random encounters. During mate search, theory predicts that the optimal search pattern depends on the expected distance to potential partners. A key question is whether actual males and females update their mate search patterns to increase encounter probability when conditions change. Here we show that two termite species, *Reticulitermes speratus* and *Coptoterines formosanus*, adaptively alternate between sexually monomorphic and dimorphic movements during mate search. After leaving their nests in a synchronized manner, termites begin to search for a mate. The resulting pairs perform tandem runs toward potential nest sites. We found that both sexes moved faster and in straight lines before finding partners, which is known to improve encounter rates when targets have completely unpredictable positions. In stark contrast, when pairs were accidentally separated during tandem running, they showed distinct sexually dimorphic movements, where females paused for long periods while males paused only briefly and moved actively. Data-based simulations demonstrated that such sexually dimorphic movements are advantageous when a mate is located nearby but its exact location is unknown. These results emphasize the importance of biological details to evaluate the efficiency of random search in animals. By extending the concept of mutual search beyond the context of mating, the dimorphic movements between partners represent a remarkable convergence between termites and other animals including humans.

**Significance Statement:** How should females and males move to search for partners whose exact location is unknown? Theory predicts that the answer depends on what they know about where targets can be found, indicating that the question doesn’t make sense until the searching context is clarified. We demonstrated that termites adaptively switch their search modes depending on the potential distance to their partners. When the location of potential mates was completely unpredictable, both sexes moved in straight lines to explore widely. In contrast, when the stray partner was at least nearby, males moved while females paused. Simulations confirmed that these movements increase the rate of successful encounters. The context-dependent switch of search modes is a key to enhance random encounters in animals.

## Introduction

Moving for survival and reproduction is nearly a defining characteristic of animals (1, 2). Their movement patterns to find targets depend on the availability of sensory information and/or memories about targets (3, 4). When searching for targets with little or no locational information, animals adopt random walks (5, 6). A number of theoretical studies have collectively revealed that the optimal movement patterns for random walks depend on the searching conditions, including density, distribution and movement of targets (3, 4, 7–18). Therefore, the detailed conditions must be specified to accurately evaluate the efficiency of movements observed in animals (4, 19). Some studies have tried to test how animals adaptively search for targets by focusing on dramatic changes of search modes depending on environmental conditions (20–23). However, as the search contexts are rarely exactly specified, most predictions remain empirically untested.

Mating is one of the main motivations of search. Optimal mate search movements are sometimes sexually dimorphic as a result of mutual evolutionary optimization (24–27). In monogamous mating systems, in which each individual mates with only one partner, theory predicts that the degree of sexual dimorphism in search strategies should depend on the potential distance to a partner, if partners are completely randomly positioned, both males and females are expected to move in a straight line, but if there is a specific available search time and expected distance to a (potential) partner, then the optimal movement patterns can be sexually dimorphic (26). To test this prediction, we focused on the mating biology of subterranean termites (Rhinotermitidae). These termites face the challenge of finding a mating partner under a variety of uncertain conditions. During a brief period of the year, alates (winged adults) fly out of their nests and disperse by wind (Fig. 1). After dispersing, both males and females land on the ground, shed their wings and run to search for a mating partner (28, 29) (Fig. 1). In this process, dealates (wing-shed adults) should have no idea of the surrounding environment because they come out of their nests for the first time. Moreover, their mate search behavior is poorly informed because the weak pairing pheromones emitted by females are effective only within a few centimeters or on contact (30). Upon joining successfully, a pair performs tandem running in which the male follows the female by maintaining almost constant contact with her back in a highly coordinated manner (31) (Fig. 1). They seek a suitable site to establish their nest, where they form a lifelong monogamous royal pair.

There are two situations where females and males have to manage uncertainty about the location of their partner during the mating process. First is when they completely lack locational information of a potential partner before pair-formation. Second is when the pair is accidentally separated after pair-formation (28). In the latter case, they know that the partner must be at least nearby, although the exact location is uncertain (Fig.1; see also *SI* text and Fig. S1). We hypothesized that termite dealates alter their movement patterns depending on the two situations described above, and thereby promote the efficiency of partner search.

**Fig. 1.**
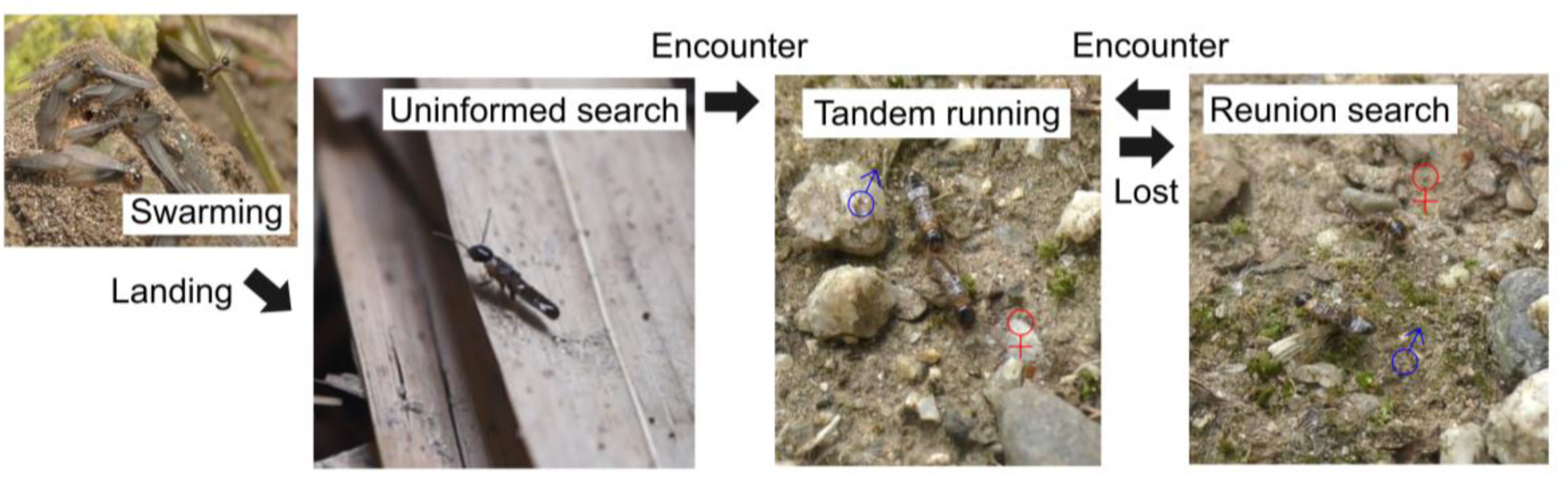
Mating biology of subterranean termites, illustrated by *Reticulitermes speratus.* In the mating season, alates (winged adults) fly off in large swarms to disperse. After dispersing, individuals shed their wings and walk to search for a mating partner without any prior locational information. Successfully encountered pairs then run in tandem to search for a suitable nest site, with the male following the female. Tandem running is quite synchronized, where a pair moves like a single individual. Nevertheless, they are sometimes separated and must search to find each other again. Thus, there are two search situations with different uncertainty about the location of their partner (before pair-fommation: uninformed search; after separation: reunion search).

We investigated the movement patterns of dealates of two termite species, *Reticulitermes speratus* and *Coptotermes formosanus*, focusing on sex differences during mate search. Both of these species show a typical sequence of mate search and monogamous colony foundation. We first observed the free-walking behavior of dealates before pair-formation, and then added a mating partner to induce tandem running. Then we removed the mating partner from the tandem to observe the change in their searching behavior after separation (Movie S1 and S2). Their motion was characterized by movement speed, turning pattern, and behavioral intermittency (pause-move patterns) as the main elements of a random walk (16). Using parameter values estimated from observations, we simulated their movement patterns under the respective searching situations to evaluate their efficiency.

## Results

### Quantitative analysis of partner search by termites

Overall, the movement patterns of both *R. speratus* and *C. formosanus* were sexually monomorphic before pair-formation as well as during tandem running after pair-formation, whereas they showed remarkable sexual dimorphism when separated after pair-formation (Fig 2). Before pair-formation, search was characterized by active movements irrespective of sex. The standard deviations of movement components overlapped between sexes, including pause duration (Fig. 2*B*, *C*), movement speed (Fig. S2) and turning angle (Fig. S3) (see *Materials and Methods).* During tandem running, movements of the dealates were also sexually monomorphic but with small differences among pairs (Fig. 2, S2 and S3). In stark contrast, the termites showed distinct sexually dimorphic movements immediately after separation: females paused for long periods while males paused only briefly and moved actively (Fig. 2; Movie S1 and S2). Over time, their movements gradually returned to the state before pair-formation (Fig. 2, S2 and S3). This sexual dimorphism was observed for a longer period in *R. speratus* (Fig. 2*B*) than in *C. formosanus* (Fig. 2*C*). *R*. *speratus* also showed a sexual difference in the moving speed (Fig. S2*A*) but not in the turning angle (Fig. S3*A*), while *C. formosanus* showed sexual differences in both (Fig. S2*B* and S3*B*).

**Fig. 2.**
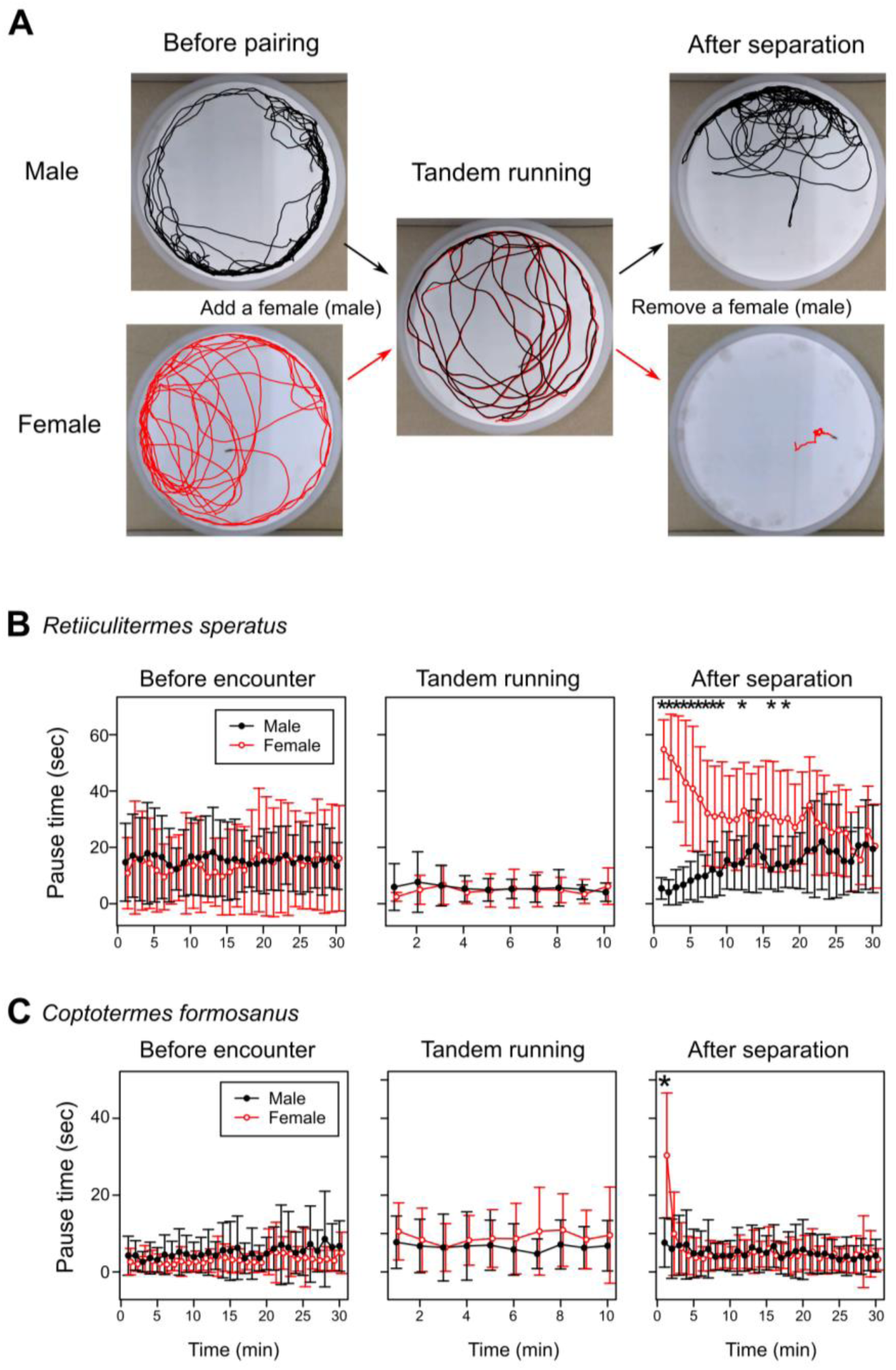
Movement patterns of termite dealates across different periods of mate search. We first observed free-walking behaviors before pair-formation. Then we added a single mating partner of the other sex to observe their movements during tandem running. Finally we carefully removed the partner using an aspirator and again observed free-walking behaviors. (*A*) Representative trajectories of dealates of *Reticulitermes speratus.* Trajectories are for 3 minutes before pair-formation, during tandem running and after separation. Before pair-formation, males and females showed similar movements, whereas they showed distinct sexually dimorphic movements after separation (Movie S1). (*B, C*) Comparison of pause time between sexes in (B) *R*. *speratus* and (C) *Coptotermes formosanus.* Both species showed sexually dimorphic movements immediately after separation, where females often paused and males moved actively. Points with bars represent mean values with standard deviations. * indicates significant differences between sexes (Wilcoxon rank-sum test with Bonferroni corrections, *P*< 0.05/70).

To evaluate the searching efficiency of the observed behaviors, we parameterized their movement patterns by decomposing them into movement speeds, turning patterns and pause-move patterns. As sexual dimorphism was the most prominent during the first minute after separation, we extracted parameters of movement patterns from data covering 1 minute each before pair-formation and after separation in *R. speratus* and *C. formosanus.* We assumed that the movement patterns of termites follow a correlated random walk (CRW), which is often used to model insect movements (11, 32) and is described by two parameters: the speed and the sinuosity (see *Materials and Methods).* Table 1 summarizes the parameter estimates. Before pair-formation, there were no statistically significant sexual differences for speed and sinuosity in either species (Table 1). After separation, we detected significant sexual differences for speed in *R. speratus* (Table 1A), and for both speed and sinuosity in *C. formosanus* (Table 1B). The changes of parameter values were observed in both sexes. Both males and females shifted to more sinuous CRWs after separation, but females also slowed their movements (Table 1).

**Table 1.** Parameters extracted from movement patterns of termites and comparison between sexes and searching schemes (before pair-formation and after separation).

**A** *Reticulitermes speratus*
| Speed | | Mean ± S.E. (mm/sec) | | Statistics | | Sinuosity | | Mean ± S.E. | | Statistics | |
| --- | --- | --- | --- | --- | --- | --- | --- | --- | --- | --- | --- |
|  |  | M (19) | F (14, 16) | W | P |  |  | M (19) | F (13, 16) | W | P |
| **Scheme** | Before | 14.7 ± 0.6 | 14.8 ± 0.7 | 154 | 0.961 | Before | | 0.85 ± 0.01 | 0.86 ± 0.01 | 189 | 0.385 |
| | After | 15.0 ± 0.8 | 5.4 ± 0.3 | 0 | **< 0.0001** | After | | 0.77 ± 0.01 | 0.77 ± 0.06 | 154 | 0.461 |
| **Statistics** | V | 106 | 0 |  |  | V |  | 9 | 29 |  |  |
|  | P | 0.679 | **0.0001** |  |  | P |  | **0.0001** | 0.2734 |  |  |

**B** *Coptotermes formosanus*
| Speed | | Mean ± S.E. (mm/sec) | | Statistics | | Sinuosity | | Mean ± S.E. | | Statistics | |
| --- | --- | --- | --- | --- | --- | --- | --- | --- | --- | --- | --- |
|  |  | M (22) | F (20) | W | P |  |  | M (22) | F (20) | W | P |
| **Scheme** | Before | 20.7 ± 1.4 | 22.3 ± 0.9 | 279 | 0.142 | Before | | 0.84 ± 0.01 | 0.85 ± 0.01 | 274 | 0.18 |
| | After | 19.0 ± 1.2 | 10.1 ± 1.1 | 43 | **< 0.0001** | After | | 0.82 ± 0.01 | 0.85 ± 0.01 | 310 | **0.0231** |
| **Statistics** | V | 104 | 0 |  |  | V |  | 15 | 35 |  |  |
|  | P | 0.4826 | **< 0.0001** |  |  | P |  | **< 0.0001** | **0.007** |  |  |

The sinuosity corresponds to the scale parameter of wrapped Cauchy distributions, covering from 0 (most sinuous) to 1 (straight motion). In statistics, W corresponds to Wilcoxon rank-sum test and V corresponds to Wilcoxon signed-rank test. M and F indicate male and female, where the numbers in parentheses indicate sample size. The sample size of females in *Reticulitermes speratus* is variable between before pair-formation and after separation because some individuals only paused during analyzed duration after separation but moved before pair-formation.

In both species, females paused for more than a half of the period (Fig. S4). We analyzed move-pause patterns of females after separation (see *Materials and Methods)*, because the move-pause intermittency is considered to be an elemental component of movement patterns (16, 33); the durations of pauses and moves often follow power-law distributions (34, 35). When we fitted power-law and exponential distributions to the data, maximum-likelihood estimations and Akaike weights indicated that both pause and move durations decayed slower than expected from exponential decay and were well approximated by power-law distributions (Fig. S5). The estimated scaling exponents were 2.675 for move duration and 1.577 for pause duration in *R. speratus;* 1.851 for move and 2.019 for pause in *C*. *formosanus.*

### Assessment of searching efficiency of random walks by termites

We used simulations to test the searching efficiency of termites’ movement patterns under respective search situations (see *Materials and Methods).* The mate search before pair-formation is the “uninformed search,” where multiple females and males search for mating partners and distances among them are completely unknown (Fig. 3*A*). We modelled this situation with two-dimensional periodic boundary conditions (size = *L* × *L;* Fig. 3*A*). Periodic boundary conditions can easily approximate infinite space with infinite number of individuals, and are often used in models or simulations of random search problems (3, 12, 13). The size *L* was set as 223.6 from the population density required for dealates of *R. speratus* to succeed in mating (20 pairs / m^2^) (36). On the other hand, the mate search after separation is the “reunion search,” where a female and a male search for their specific partner nearby without detailed locational information (Fig. 3*D*). This situation was modelled by a borderless continuous space, where a female and a male were initially separated by the distance *d* (Fig. 3*D*). To obtain the distance *d*, we observed spontaneous separations during tandem running and measured the separated distance between a male and a female every 0.2 seconds until they re-encountered (see *SI text).* The distance *d* was determined to be 16.09 mm for *R. speratus* and 22.97 mm for *C. formosanus* respectively as the mode value of the distribution of separated distance. We evaluated four possible combinations of movement patterns: the observed sexually dimorphic movement after separation, the observed sexually monomorphic movement before pair-formation, a virtual monomorphic movement where both partners moved like females after separation, and a virtual monomorphic movement were both moved like males after separation.

**Fig. 3.**
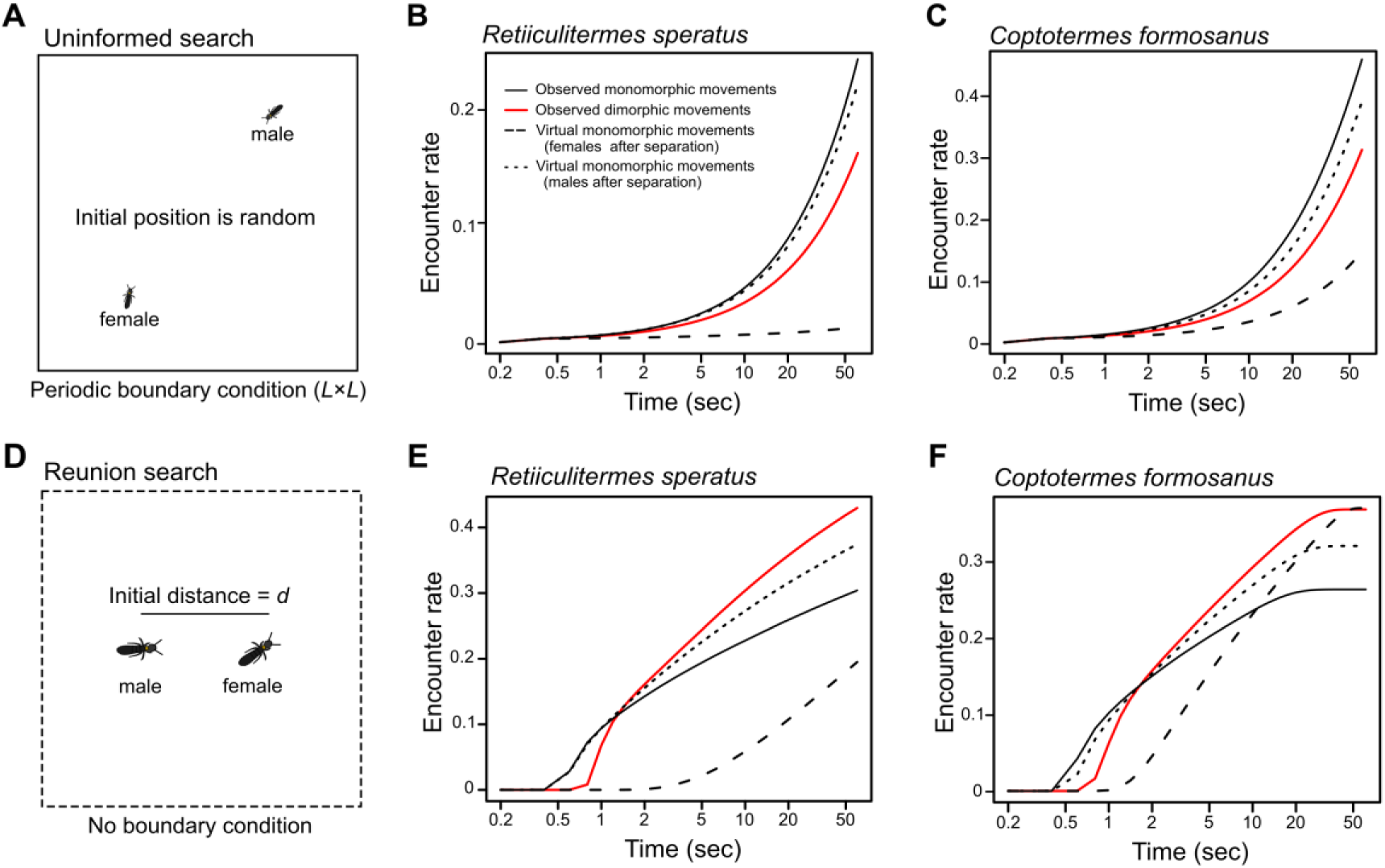
Searching efficiency of termite movement patterns in two different searching situations. *(A)* Assumed search conditions before pair-formation (uninformed search). Before pair-formation, females and males search for partners without any locational information. Such conditions are simulated by periodic boundary conditions (size *L* × *L)* with random initial positions of females and males. (*B, C*) Searching efficiency of observed movement patterns under uninformed search conditions for (*B*) *Reticulitermes speratus* (*L* = 223.6, φ = 7) and (*C*) *Coptotefmesformasanus* (*L* = 223.6, φ = 10). In both species, sexually monomorphic movements observed before pair-formation achieved higher encounter rates than the other combinations of movements. (*D*) Assumed search conditions after a tandem running pair gets separated (reunion search). After separation, the female and the male search for each other, where they cannot know the detailed location of the partner although the distance between them is expected to be short Such conditions am simulated by borderless continuous spaces with a distance *d* between a female and a male. (*E, F*) Searching efficiency of observed movement patterns under reunion search conditions for (E) *R. speratus (d* =16.09, φ = 7) and (*F*) *C. formosanus* (*d* = 22.97, φ = 10). In turn, sexually dimorphic movements observed after separation achieved higher encounter rates than the other sexually monomorphic movements during realistic searching periods in both species. The movements of individuals are modeled by correlated random walk with or without pauses, where parameters are shown in Table 1. The results were obtained from means of 1,000,000 simulations.

Under the uninformed search conditions, the observed sexually monomorphic movement always achieved higher efficiency than the observed sexually dimorphic movement or the other virtual monomorphic combinations (*R. speratus:* Fig. 3*B*. *formosanus:* Fig. 3*C*). Under the reunion search conditions, the results depended on the length of search time (Fig. 3*E*, 3*F*), which had been predicted in the previous theoretical study (26). The sexually dimorphic movement achieved higher efficiency than the sexually monomorphic combinations when the search time was longer than ca. 2 sec (and shorter than 47.4 sec in *C. formosanus)*, while some sexually monomorphic combinations achieved higher encounter successes outside these ranges (Fig. 3*E*, *F*). Based on our empirical observations, 100% and 85% of reunion searches for *R. speratus* (*n* = 282) and *C. formosanus* (*n* = 56), respectively, ended with re-encounter within the above ranges (Fig. S1; *SI text*), illustrating that the sexually dimorphic movement was the best way for the termites to facilitate reunion in realistic search time.

## Discussion

Our simulations parameterized by empirical behavioral measurements demonstrated that the transition of different movement modes observed in termites is adaptive in promoting encounters across different informational contexts (Fig. 3), as follows. First, before pair-formation, individuals have no information to locate their potential partners, and they engage actively in random search with relatively straight motions (i.e., higher ρ values; Table 1). This should be adaptive, compared to their alternative movements, because the optimal strategy is moving straight without directional changes when targets are destructive (i.e., disappear after mating) and randomly distributed (3, 15, 26). Second, when a mated pair gets separated after pair-formation, the termites generally increase their turning rate and show remarkable sexually dimorphic movements: females pause frequently and males move actively (Fig. 2). They have information that their mating partner must be nearby, and our simulations demonstrate that their movements are adaptive responses to reunion search (Fig. 3). This dimorphism in pausing behavior indeed facilitates reunion: without pausing by females, the pair would suffer from lowered encounter efficiency *(SI text* and Fig. S6). Finally, over 30 minutes of separation, the termites progressively change their movements from sexually dimorphic to monomorphic ones (Fig. 2). This suggests that this duration of searching in vain is enough for the termites to decide that the lost partner is no longer near them. Our results support the notion that animals do not engage in a one-size-fits-all search strategy but update their strategies through feedback from their experience (4, 20, 37, 38).

A comparison between our mate search system and prey-predator systems gives an insight into flexibility of animal movements. The contrasting movements between random and reunion searches are reminiscent of previous observations that predators often alternate between extensive and intensive searches depending on prey distributions (2, 4, 21, 22). In extensive searches, predators explore by moving for longer distances to find more profitable areas, whereas in intensive searches, they exploit prey-rich areas with frequent turns. Moreover, during reunion searches, females engaged in pausing to facilitate encounters. Similar animals which show pausing behavior in biological encounters are ambush predators, where the advantage of pausing or moving should depend on various factors such as body size and the abundance of targets (35, 39). By focusing on when pausing behavior offers advantages, our results shed light on the reverse side of random search problem that asks for the most efficient way to be found by specific moving agents.

In the model of reunion search, the sex roles should be symmetric and interchangeable. Then, why was the pausing sex always females in termites? A reasonable explanation invokes the sexual difference in detection ranges. Tandem running of termites is often mediated by pair-bonding pheromones secreted by leading females to help males follow them (30). This pheromone should enable males to detect females from a longer distance than females. A previous model predicted that, with such asymmetric detection ranges, high efficiency of encounters can be achieved when the sex with a narrower detection range moves slower or even pauses (27). Moreover, *R. speratus* showed a larger sexual difference than *C*. *formosanus*, which also supports the previous model which predicts that the “short-sighted” sex should move more slowly under larger sexual differences in the detection range (27). The pair-bonding pheromone of *Reticulitermes* termites is volatile (40, 41) whereas that of *C. formosanus* is non-volatile and works upon contact (42). Therefore, *R. speratus* males can detect partner females from a longer distance than *C*. *formosanus* males. It should be noted that we did not observe sexually dimorphic movements before pair-formation, suggesting that the mere presence of sexual difference in the detection range due to attracting signals cannot fully explain the movement patterns of termites. Integrating the effect of attracting signals and the optimal random search enabled a fuller understanding of the flexible random search strategies of termites.

An important goal in the study of optimal random search is to identify the common features of the underlying processes across species (43). The advantage of dimorphic movements in reunion search revealed in this study can be applied to various social interactions beyond the mate search context. For example, ants perform tandem running during nest relocation and group foraging, in which a single well-informed worker guides a naïve nestmate to a target site (44, 45). When the follower loses its leader, the leader pauses and waits for the missing follower, while the follower engages in a Brownian walk that is effective to search for the leader (46). Although form and function of tandem running are different between ants and termites, our results represent a remarkable evolutionary convergence of behavioral dimorphism that promotes encounter with a separated partner. The convergence should further be extended to our own species by asking how two people can find each other efficiently in a known search region like the High Street or a shopping mall, known as the rendezvous search problem in operations research (47–49). This problem has many variations, and it can sometimes be optimal for one player to wait for the other (50). Our findings thus have potential applications to the design of human-engineered searchers (43). Combined with the previous studies, our results illustrate the convergence between adaptive evolution of animals and rational thinking of humans.

To conclude, biological details are essential when we evaluate the random search strategies of animals. Types of targets (9, 26, 51) and searching conditions (8) strongly affect the search efficiency and the resulting fitness. Comparative experiments across conditions as well as across species will offer an ideal opportunity to understand how animals search for uncertain targets in complex environments. In this study, we focused on the walking behavior of termite dealates which has a sole function of mate search. Our data-driven modelling successfully demonstrated that the termites adaptively switch between sexually monomorphic and dimorphic movements depending on the informational contexts, which improved pairing compared to the alternative strategies. It is usually difficult for animals to achieve optimal states in a strict sense that are theoretically derived (52). Nevertheless, the condition-dependent changes along continuous parameters clearly indicate the adaptive value of respective search strategies. A comprehensive view of adaptive search strategies that is achieved by clarifying their contexts will have wide applications that should contribute to our well-being, as well as deepen our understanding of life.

## Materials and Methods

### Termites and experimental setup

We used two termite species, *Reticulitermes speratus* and *Coptotermes formosanus,* which show a common sequence of mating behavior: they walk to search for mates and the resulting pairs perform tandem running to seek potential nest sites (29). We collected alates of *R*. *speratus* from five colonies (R_A_–R_E_) together with a piece of nesting wood from oak or Japanese cedar forests in Kyoto, Japan in May, 2017, just before their swarming season. Alates of *C. formosanus* were collected using light trapping (79 individuals, C_L_) as well as from nesting wood from pine forests (two colonies, C_A_ and C_B_) in Wakayama, Japan in June 2017. All collected individuals were maintained in artificial nests at 20 °C under dark conditions until the experiments to control flight timing. Just before each experiment, we transferred the nests into a room at 27 °C, which promoted alates to emerge and fly. Alates were then collected and separated individually. We used individuals that shed their wings by themselves within *24* hours.

To observe mate search behavior of termite dealates, we prepared an experimental arena by filling a petri dish (φ = 145 mm) with moistened plaster, so that the surface of the arena can be cleaned by slicing off plaster before each trial. A video camera (BSW50KM02BK USB2.0 camera, BUFFALO, Japan) was mounted vertically above the arena and the camera system adjusted so that the arena filled the camera frame. The video was recorded to a Windows PC using CCI-Pro-MR (http://www.cosmosoft.org/CCI-Pro-Mr/) at a resolution of 640 × 480 pixels. We extracted the coordinates of termite movements at a rate of 5 times per seconds from each video using the video-tracking system UMATracker (http://ymnk13.github.io/UMATracker/). The arenas were illuminated by two white LED lights with an intensity of 430 ± 60 lux. In the observation of termite behavior, each individual was used only once. All data analyses were performed using R v3.1.3 (53) and all data are available in *SI*.

### Analysis on mate search movements of termites

We analyzed movement patterns of termite dealates from before pair-formation to after separation. We placed a single dealate on the experimental arena and observed its free-walking behavior for 30 minutes. Then we added a single mating partner of the other sex. If they formed a tandem within 5 minutes, we observed their tandem running behavior for 10 minutes. After 10 minutes, we carefully removed the added partner using an aspirator and again observed free-walking behavior for 30 minutes (Movie S1, S2). In case a tandem pair was not formed within 5 minutes, we ceased the trial. We obtained full behavioral observations of 19 males (R_A_: 5, R_C_: 4, R_D_: 6, R_E_: 4) and 18 females (R_A_: 5, R_C_: 3, R_D_: 6, R_E_: 4) in *R. speratus*, and 23 males (C_A_: 7, C_B_: 4, C_L_: 12) and 23 females (C_A_: 8, C_B_: 3, C_L_: 12) in *C. formosanus.*

We first compared the movement patterns between sexes. We described the movement patterns of termites by move-pause patterns, moving speed and turning patterns. First, the movements of each individual were discretized into a series of moves and pauses. The distribution of the length of displacements between successive frames (0.2 s) was bimodal with two peaks around 0 mm and 3.5 mm for *R. speratus*, and 0 mm and 4.7 mm for *C. formosanus.* The two peaks can be regarded as representing pauses and moves, respectively. We obtained the thresholds for move/pause as the value representing the second peak multiplied by 0.2 (= 0.70 mm for *R. speratus;* 0.94 mm for *C. formosanus)*, where pause was defined as displacement less than or equal to the thresholds (the positions of the values in the histograms were shown in Fig. S7). These values were close to our visual discrimination of move and pause during video observations. We examined other threshold values, but it did not affect our conclusions qualitatively. We calculated the duration of pauses in every minute for before encounters (30 minutes), during tandem running (10 minutes) and after separation (30 minutes). The moving speeds were computed by the instantaneous speed when termites were moving. The turning angles were computed as the magnitude of changes in the direction of motion from one frame to the next frame during moving. We also calculated turning angles for the direction changes before and after pauses. Then the mean value of moving speeds and turning angles were obtained in every minute for before pair-formation (30 minutes), during tandem running (10 minutes) and after separation (30 minutes). These three components (duration of pauses, moving speeds and turning angles) were compared between sexes every minute using Wilcoxon rank sum test with Bonferroni correction for multiple comparison (α = 0.05/70).

As a result of the above analysis, we found that the sexual dimorphism was prominent just after separation in both species (Fig. 2*B, C*). We focused on 1 minute after separation to explore the difference in movement patterns between before pair-formation and after separation. For comparison, we prepared those of the last 1 minute before pair-formation. We modelled movement patterns of termites by correlated random walks (CRW), which accounts for the emergence of angular correlations in animal trajectories coming from local scanning behavior (16, 32, 54). CRW can be described by two parameters: speed and sinuosity. The speed was obtained by the mean of moving speeds. The sinuosity was given by the scale parameter which was obtained by fitting turning angle data to wrapped Cauchy distributions using maximum likelihood estimation methods. Wrapped Cauchy distribution covers from uniform distribution (scale parameter = 0) to delta distribution (scale parameter = 1) and corresponding movement patterns of Brownian and straight walks, respectively (3). We tested the difference of these two parameters between before pair-formation and after separation using Wilcoxon signed-rank test, and between sexes using Wilcoxon rank-sum test.

In addition, as the pause behavior was observed especially in females after separation, we examined the frequency distribution of moving and pausing duration for them. In measuring the duration of move and pause, we observed females until the end of the behavior that was performed by them at 1 minute from the onset (i.e., the observed duration was longer than 1 min). We fitted exponential distribution and power-law distribution with a minimum value of 0.2 to all data using maximum likelihood methods (55) with model selection using the Akaike weight.

### Simulations

We developed an individual based model to examine the efficiency of movement patterns observed in termites. We prepared two searching situations, a periodic boundary condition of size = *L* × *L*, and a borderless continuous space where a female and a male initially separated by the distance *d.* The former was to simulate uninformed search before pair-formation and the latter was to simulate reunion search after separation. In each condition, we considered a female and a male walking until encountering another individual of the other sex. When the distance between the centers of the two individuals became smaller than φ, they were regarded to encounter. This φ value is based on the definition of tandem running, where 7 mm for *R. speratus* and 10 mm for *C. formosanus* (see *SI* text).

Individuals perform CRW with the parameters of speed and sinuosity, denoted by *v* and ρ, respectively. These parameters were obtained as the mean value of the observation data for each sex and search condition (Table 1). Based on our behavioral analysis, each time step was adjusted to 0.2 seconds. Thus, each individual moved 0.2v mm in each time step. Turning angles followed wrapped Cauchy distribution with scale parameter ρ. After generating a uniform random number *u* (0 < *u* ≤ 1), the turning angles θ were derived from the following equation by applying the inversion method (16):

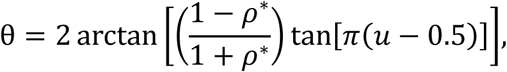

where we did not include the median parameter because animals performing CRW maintain the previous direction while scanning (16). For the initial conditions, we assumed that moving directions of all individuals were random.

We added intermittency to the above CRWs for females after separation to account for pausing behaviors. Observations showed that their distributions of the duration of move and pause followed power-law distributions. Thus we considered that females after separation repeated move for tm and pause for *t*_p_. After generating a uniform random number *u* (0 < *u* ≤ 1), the duration *t*_m_ and *t*_p_ were derived from the following equation: *t*_rn_ or *t*_p_ = *t*_0_*u*^1^/^(1–μ)^, where *t*_0_ is 0.2 seconds and μ is the power-law exponent. The values of μ were 2.675 for move duration and 1.577 for pause duration in *R. speratus*; 1.851 for move and 2.019 for pause in *C. formosanus.* For the initial condition, individuals were assumed to perform moving or pausing with the probability of 0.5.

We compared the searching efficiency among four possible combinations of movement patterns: the observed sexually dimorphic movement after separation, the observed sexually monomorphic movement before pair-formation, two virtual monomorphic movements with both a female and a male moving like females or males after separation, for each searching condition in each species. Simulations were performed for 60 seconds (= 300 time steps), based on the duration of empirical measurements. We ran 1,000,000 simulations and measured the efficiency as the probability to encounter a mating partner. The simulation program was implemented in Microsoft Visual Studio C++ 2017. The source codes for all simulations are available in *SI.*

## Acknowledgments

We thank Masato S Abe for helpful advice on data analysis; Stephen C Pratt for helpful comments which improved the manuscripts and English corrections; Kenji Matsuura, Naohisa Nagaya, Ryusuke Fujisawa, Chin-Cheng Scotty Yang, Hiraku Nishimori and the members of the Laboratory of Insect Ecology in Kyoto University for fruitful discussion; Osamu Yamanaka for suggestion of the use of UMATracker; Taisuke. Kanao and Aoi Mizumoto for their help with termite collection. NM completed writing the manuscript supported by JSPS Overseas Research Fellowships. This study was supported by Grants-in-Aid for JSPS Research Fellow 15J02767 (NM), 15K18609 and 17H01249 (SD)..

